# Early IKKβ-dependent anabolic signature governs vascular smooth muscle cells fate and abdominal aortic aneurysm development

**DOI:** 10.1101/2024.06.12.598763

**Authors:** Priscilla Doyon, Ozge Kizilay-Mancini, Florence Dô, David Huynh, Gaétan Mayer, Stéphanie Lehoux, Huy Ong, Maelle Batardière, Vincent Quoc-Huy, Sylvie Marleau, Simon-Pierre Gravel, Marc J Servant

**Affiliations:** Faculty of Pharmacy, Université de Montréal, Montréal, Canada; McGill University Health Centre, McGill University, Montréal, Canada; Institute for Research in Immunology and Cancer, Université de Montréal, Montréal, Canada; The Montreal Heart Institute, Université de Montréal, Montréal, Canada; Lady Davis Institute for Medical Research, McGill University, Montréal, Canada

## Abstract

**Background:** Abdominal aortic aneurysm (AAA) is a detrimental disease with no effective pharmacological therapy. While inflammation is recognized as one of the key regulators of AAA, targeting inflammatory pathways once the disease is established does not impact the outcomes. However, understanding the earliest molecular indicators could shed light on the precise biological targets and prognostic markers for AAA.

**Methods:** Using apolipoprotein E (*ApoE*)-deficient mice fed with a standard diet and infused with Angiotensin II (Ang II), we conducted bulk RNA-sequencing (RNA-Seq) analysis on suprarenal (SRA) regions obtained from both unchallenged and challenged WT mice, specifically examining responses 24 hours after Ang II infusion to capture the initial phases of aortic stress response. We further created a unique model of hyperlipidemic mice in which the expression of the inhibitor of nuclear factor kappa B kinase subunit beta (IKKβ) can be conditionally (via tamoxifen injection) suppressed in vascular smooth muscle cells (VSMC). The development of AAA was evaluated using in situ examination and quantified using RT-qPCR, immunohistochemistry and fluorescence microscopy. Cultured VSMC were exposed to the selective IKKβ inhibitor MLN120b and the expression levels of phenotypic markers kruppel like factor 4 (KLF4), β-Catenin and cellular communication network factor 2 (CCN2) were addressed using cellular extracts and immunoblot analysis.

**Results:** RNA-Seq data support the presence of early anabolic events in SRA regions detailing activation of the mammalian target of rapamycin complex 1 (mTORC1) pathway, which paralleled cellular anabolic processes including mitochondria, ribosome and sterol biosynthesis, the Unfolded Protein Response (UPR) and fibrogenesis. Conditional deletion of the *Ikbkb* gene in VSMC significantly reduces the incidence of SRA lesions as well as the rate of aneurysm ruptures in mice exposed to Ang II. *In situ* analysis further demonstrated that the protection conferred by the lack of IKKβ expression in VSMC is associated with reduced inflammatory response, the preservation of the contractile over the degradative VSMC phenotypes, and the absence of an anabolic signature.

**Conclusion:** Our results not only reinforce the major roles played by VSMC in the rapid adaptation leading to the deleterious remodeling of the vascular wall and aortic lesions but also support a paradigm aiming at repositioning the efforts focusing on anabolic rather than inflammatory events.

## INTRODUCTION

Abdominal aortic aneurysm (AAA) is a life-threatening disease defined by progressive aortic dilation caused by the imbalance between synthesis and degradation of extracellular matrix (ECM) components in the vascular wall ^1^. Clinically defined as a focal dilatation of the infrarenal aorta (> 3cm), AAA is the most common form of aneurysm in humans. Although increased leukocytic infiltration and a decreased number of vascular smooth muscle cells (VSMC) in the adventitial and medial layer correlates with the extent of expansive remodeling (i.e. aortic dilatation) and consequently aortic dissections and rupture ^2^, the precise mechanisms underlying uncontrolled and exaggerated expansive remodeling remain elusive. Therefore, there is no efficient pharmacological therapy to stabilize or reverse the evolution of small AAA, with open or endovascular surgeries remaining the only option to treat this detrimental disease ^3^.

Risk factors associated with the development of AAA are well-documented and include age, male gender, smoking, hypertension, low levels of high density lipoprotein (HDL), cholesterol, and a family history of AAA ^4^. Inflammation is implicated in many of the risk factors for developing AAA with most of the studies focusing on inflammatory cells such as macrophages, neutrophils, CD4^+^ T cells, and mast cells in ECM remodeling events ^5–9^. Another feature that is common to numerous risk factors for developing AAA is the presence of various anabolic processes ^10^; yet, this aspect is not studied as much as inflammation.

VSMC that reside in the tunica media are the major component of the aortic wall. They contribute to vascular hemostasis through their secretion of cross-linked collagen and elastin. Under normal conditions, VSMC are quiescent and non-migratory, displaying a contractile differentiated phenotype ^11, 12^. However, VSMCs are not terminally differentiated and undergo reversible changes (or dedifferentiation) towards a so-called « synthetic » phenotype, a process referred to as phenotypic modulation, cell plasticity, or switching, in response to vascular injuries. VSMC phenotypic switching is characterized by reduced expression of SMC-selective differentiation markers (smooth muscle protein 22-α (SM22A), α-smooth muscle actin (ACTA2)) and increased VSMC growth (hypertrophy and/or hyperplasia), migration, senescence, increased ECM metabolism, reactive oxygen species (ROS) production, and inflammatory responses ^13^. Interestingly, this somehow “inflammatory/hyperactive-synthetic state” of VSMC was first observed and characterized in human and animal models following angioplasty, in hypertension-associated remodeling events in small arteries as well as atherosclerosis^12,14,15^. Importantly, it has more recently been described in humans and mice suffering from AAA ^16–18^. However, the role of VSMC-dependent anabolic pathways in AAA development and subsequent rupture and dissection remains unclear.

We previously demonstrated that the inhibitor of nuclear factor kappa B kinase subunit beta (IKKβ) is expressed in VSMC and plays key roles in the inflammatory signaling events in response to vasoactive peptides and pathogens ^19–22^. Through its ability to control the activation of nuclear factor kappa B (NF-κB) transcription factors ^23^ and the stabilization of mRNAs encoding for numerous proinflammatory cytokines ^24,25^ IKKβ is a major effector of the innate and adaptive immune systems. It is therefore considered as a druggable anti-inflammatory target for cancer treatment ^26^. Although previous mouse models of AAA have shown that anti-inflammatory strategies can effectively attenuate aneurysm formation ^27–30^, quenching vascular inflammation using classical pharmacological approaches (e.g. indomethacin) or inhibiting the immune system (e.g. mast cell inhibition, anti-interleukin-1β therapy) does not influence the progression of AAA in humans ^31^. Thus far, the apparent failure in the identification of a druggable target to treat AAA could be likely explained by the inability of the anti-inflammatory approaches to target the detrimental synthetic state of VSMC that occurs during AAA development ^17^ or the lack of a complete picture of the molecular events that drive this aortopathy.

In this study we used the well-characterized murine model of AAA consisting of Angiotensin II (Ang II) infused mice on an hyperlipidemic apolipoprotein E^-/-^ (ApoE^-/-^) genetic background to demonstrate the establishment of rapid and unique anabolic molecular signature that is orchestrated by IKKβ. Furthermore, preventing the expression of IKKβ specifically in VSMC reduced the occurrence of the synthetic phenotype and attenuated the AAA formation. Our data suggest that the control of the anabolic response through IKKβ could be exploited for the treatment of AAA.

## METHODS

### Data Availability

The data that support the findings of this study are available from the corresponding author upon reasonable request. The RNA sequencing (RNA-Seq) data generated in this study have been deposited in NCBI GEO (accession number: GSE265897; key: ijkdsqasrdkdnyj). Full methods are provided in the Supplemental Material.

## RESULTS

### Early anabolic response defines the molecular signature of aortopathy in the classical Ang II-infused *ApoE*-deficient mice-AAA model

Tissue remodeling leading to aortic dissection occurs very rapidly in 8-10 weeks old male *ApoE*-deficient mice fed with a standard diet and infused with Ang II ^32,33^ and in fact, a vast majority of fatalities due to aortic ruptures occured before the first seven days of Ang II infusion and correlate with the presence of phagocytic markers in the AAA lesions, concomitant to the disruption of elastin fibers ^34^. Therefore, most studies focusing on the molecular etiology of AAA employ whole aortas or the suprarenal (SRA) regions isolated from animals exposed to Ang II for 2 to 7 days ^35^ where vascular wall remodeling (mostly ECM metabolism) associated with wide-ranging increases in inflammatory genes are already present, a situation that could misguide and complexify the comprehension of early cellular, molecular and biochemical responses underlying this aortopathy. We therefore conducted bulk RNA-Seq analysis on SRA regions obtained from both unchallenged and challenged IKKβ^+/+^ mice, specifically examining responses 24 hours after infusion with Ang II at a rate of 1000 ng/min/kg to capture the early initial phases of the aortic stress response. Volcano plot of the identified 1917 differentially expressed genes (DEG) revealed unique patterns in aortic gene expression between unchallenged and challenged IKKβ^+/+^ mice (**Figure 1A-B**). Next, we performed gene functional analyses on these differentially expressed transcripts (DET) to characterize the transcriptomic signature associated with early Ang II signaling in SRA. First, functional analysis of significantly induced DET using the Gene Ontology Biological Process (GOBP) collection of gene sets reveals enrichment of pathways associated with cell growth, cell proliferation, extracellular matrix remodeling, angiogenesis, and sterol biosynthesis (**Figure 1C).** Performing a similar analysis but using the KEGG pathway collection of gene sets indicated enrichment pathways associated with hypertrophic disease, pro-mitogenic and inflammatory signaling, as well as terpenoid and steroid biosynthesis, both linked to cholesterol biosynthesis (**Figure 1D**). Strikingly, these two functional analyses reveal that early Ang II signaling in SRA leads to transcriptomic changes associated with long-term and chronic hypertrophic disease. To better delineate this early transcriptomic signature, we performed Gene Set Enrichment Analyses (GSEA), a computational and statistical method that does not apply cutoffs and considers all genes within a gene set ^36^. GSEA analyses confirmed the enrichment of the sterol biosynthesis gene set (**Figure 1E**). Surprisingly, out of 18 enzymes involved in cholesterol biosynthesis, 15 had fold change > 1.2 fold, and 11 were significantly upregulated (adjusted p-value < 0.1) (**Suppl. Figure S1A**). In addition to cholesterol biosynthesis, GSEA analyses revealed the enrichment of metabolic pathways that functional analyses could not detect, such as ribosome biosynthesis, mitochondrial ATP synthesis, and enrichment for mammalian target of rapamycin complex 1 (mTORC1) signaling, a major regulator of metabolism vascular muscle growth (**Figure 1E**). GSEA analyses also revealed enrichment for the unfolded protein response (UPR) and fibrosis, which suggests outstanding protein synthesis and extracellular matrix remodeling that could quickly ignite stress and pro-inflammatory responses. To help identify the transcription factors that could mediate these transcriptomic changes in response to Ang II in SRA, we examined the over-representation of transcription factor binding sites (TFBS) in the promotor of significantly induced DET. These analyses showed enrichment of TFBS for transcription factors linked to vascular remodeling, anabolic and fibrotic events, including mothers against decapentaplegic (SMADs), cAMP-responsive element-binding (CREB), ETS transcription factor ELK1 (ELK1), early growth response (EGR), MYC proto-oncogene bHLH transcription factor (MYC), X-box binding protein 1 (XBP1), activating transcription factors (ATFs), NF-κB and Krüppel-like factors (KLFs) (**Figure 1F**). These findings support early anabolic events in the initiation of AAA, mainly composed of cellular processes that were recently shown to be under the direct control of the mTORC1 pathway^10,37,38^.

**Figure 1.**
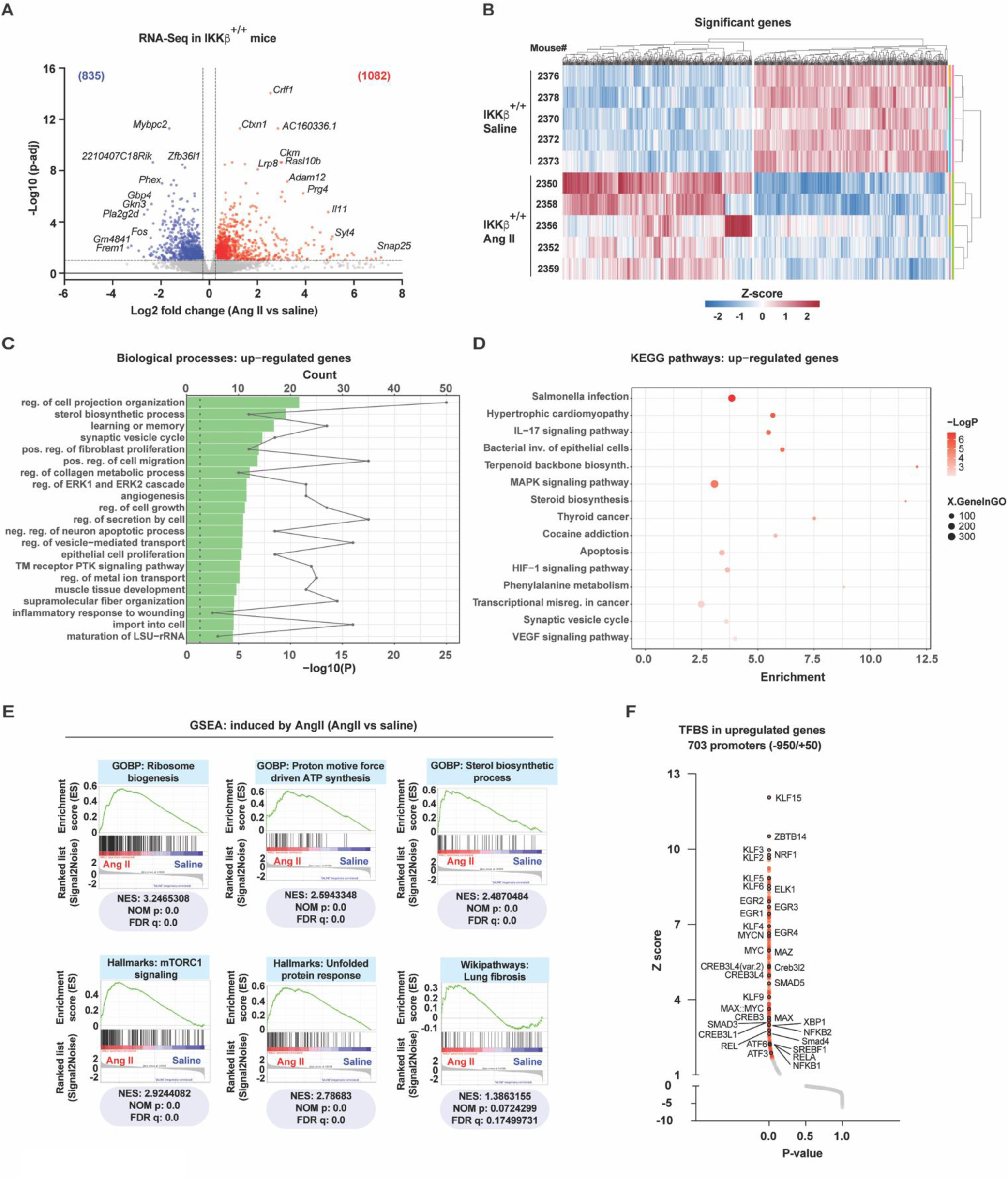
: Early anabolic response define the molecular signature of aortopathy in the classical Ang II-infused *ApoE*-deficient mice-AAA model. A) Volcano-plot representation of differentially expressed transcripts (DET) in SRA in response to Ang II treatment for 24 hours. Each dot indicates the average expression fold change and associated p-adjusted value for a single transcript when comparing Ang II (n= 5 mice) versus saline (n= 5 mice) groups. Colored numbers indicate the number of significantly induced (red) and reduced (blue) transcripts at a given threshold (dotted lines), as indicated in Materials and methods. B) Heatmap representation of significant DET. Fold changes in expression have been transformed to z-scores, and clustering has been done on rows (mice) and columns (transcripts). C) Functional classification of significant upregulated DET with Metascape for the Gene Ontology : Biological Processes collection of gene sets. Green bars indicate p-values (-Log10), and dots indicate the number of genes per pathway. D) Functional classification as in C but for KEGG pathways using enrichment bubble plot. The X-axis represents the enrichment factor (nb of genes/all genes from pathway), XgeneInGO indicates the number of genes (bubble size), and -LogP indicate significance based on the p-value. E) Gene set enrichment analyses (GSEA) done on all normalized transcripts in control (saline) and Ang II-treated mice. NES: Normalized enrichment score; NOM p: nominal p-value; FDR q : false discovery rate-adjusted q-value. F) Over-representation of transcription factor binding sites (TFBS) in the promotor (-950 to +50) of significantly induced DET in Ang II-treated versus saline-treated mice. Red dots indicate significant over-representation with a p-value < 0.05. Labelled dots also have a black outline.

### The absolute requirement of vascular wall-expressed IKKβ in pseudoaneurysm development and subsequent fatal ruptures

The inflammatory effector kinase IKKβ, known for its absolute requirement for the activation of the canonical NF-κB transcription factor pathway, was also previously shown by our group to regulate mTORC1 activation in response to Ang II, resulting in the control of both the inflammatory and hypertrophic effects of the vasoactive peptide ^21^. Given the known and widely accepted inflammatory nature of AAA and the observed anabolic mTORC1 signature above, we next questioned the role of IKKβ in the development of experimental AAA by generating hyperlipidemic mice in which the *Ikbkb* gene is specifically inactivated in VSMC. We refer to this unique transgenic homozygous hypercholesterolemic mouse model as SMA-CreER^T2^Ikkβ^Flox/Flox^*ApoE*^-/-^ where the expression of IKKβ is temporally (tamoxifen injection) and selectively (CreER^T2^ under the control of the *Acta2* promoter; refer to **Suppl. Figure 1B**) prevented in *ApoE*^-/-^ VSMC on a C57BL6 background. At 8 weeks of age, male SMA-CreER^T2^Ikkβ^Flox/Flox^*ApoE*^-/-^ (IKKβ^-/-^) and their control littermates SMA-CreER^T2^*ApoE*^-/-^ (IKKβ^+/+^) mice were subjected to 5 consecutive days of tamoxifen injection. After an additional 7 days, mice were infused using the classical AAA-induction protocol consisting of animals exposed to either saline or Ang II for 28 days a time point where vessel wall expansion, pseudoaneurysm formation, and ruptures can all be simultaneously assessed (**Figure 2A**). Tamoxifen infusion efficiently induced cell-specific conditional deletion of IKKβ in VSMC as no staining of the kinase was detected in the media compared to the intima and adventitia (**Figure 2B**). Morphologically, the aortas of saline-infused IKKβ^-/-^ animals did not differ from those of saline-infused control IKKβ^+/+^ mice (**Suppl. Figure 1C**). Moreover, IKKβ deficiency did not affect body weight, end-point systolic blood pressure, plasma cholesterol and triglycerides. (**Suppl. Figure 1D-E**). Systemic Ang II infusion at the classical dose of 1000 ng/kg/min successfully induced AAA in hypercholesterolemic mice and the deletion of IKKβ in VSMC dramatically reduced the incidence and the maximal abdominal aortic diameters of the aortic SRA lesions while increasing the survival of the animals. (**Figure 2 C-E**). Importantly, fatalities due to abdominal aortic ruptures were diminished by ∼48% in the IKKβ^-/-^ group allowing more animals from this group to develop complex and stable lesions over the 28 days of treatment (**Suppl. Figure 1F-G)**. Next, SRA segments isolated from animals that survived the 28 days treatment were further analyzed for classical endpoints of aortic wall remodeling and inflammation. Elastin degradation and adventitial macrophage recruitment were significantly reduced in the absence of IKKβ in the media, effects that corresponded to healthier medial regions composed of densely populated, multilayered SMC (**Figure 2F-H**). Furthermore, adjacent tissues composed of the end of the descending thoracic aorta (TA) and the beginning of the SRA segments (TA/SRA) were isolated from randomized animals described above and used to isolate RNA and interrogate the expression of mRNA transcripts that have been linked to aortic disease. In Ang II-treated animals, the aortas depicted an almost total dependence of the presence of IKKβ in medial VSMC for the induction of classical inflammatory mRNA transcripts composed of cytokine/chemokine (interleukin-1β (*Il1b*)/ C-C motif chemokine ligand 2 (*Ccl2*)), adhesion molecules that regulate leukocytes recruitment (vascular cell adhesion molecule 1 (*Vcam1*)/ intercellular adhesion molecule 1 (*Icam1*)) as well as ECM remodeling enzymes (matrix metallopeptidase 2 (*Mmp2*), 9 (*Mmp9*) and 14 (*Mmp14*)) (**Figure 2I**).

**Figure 2.**
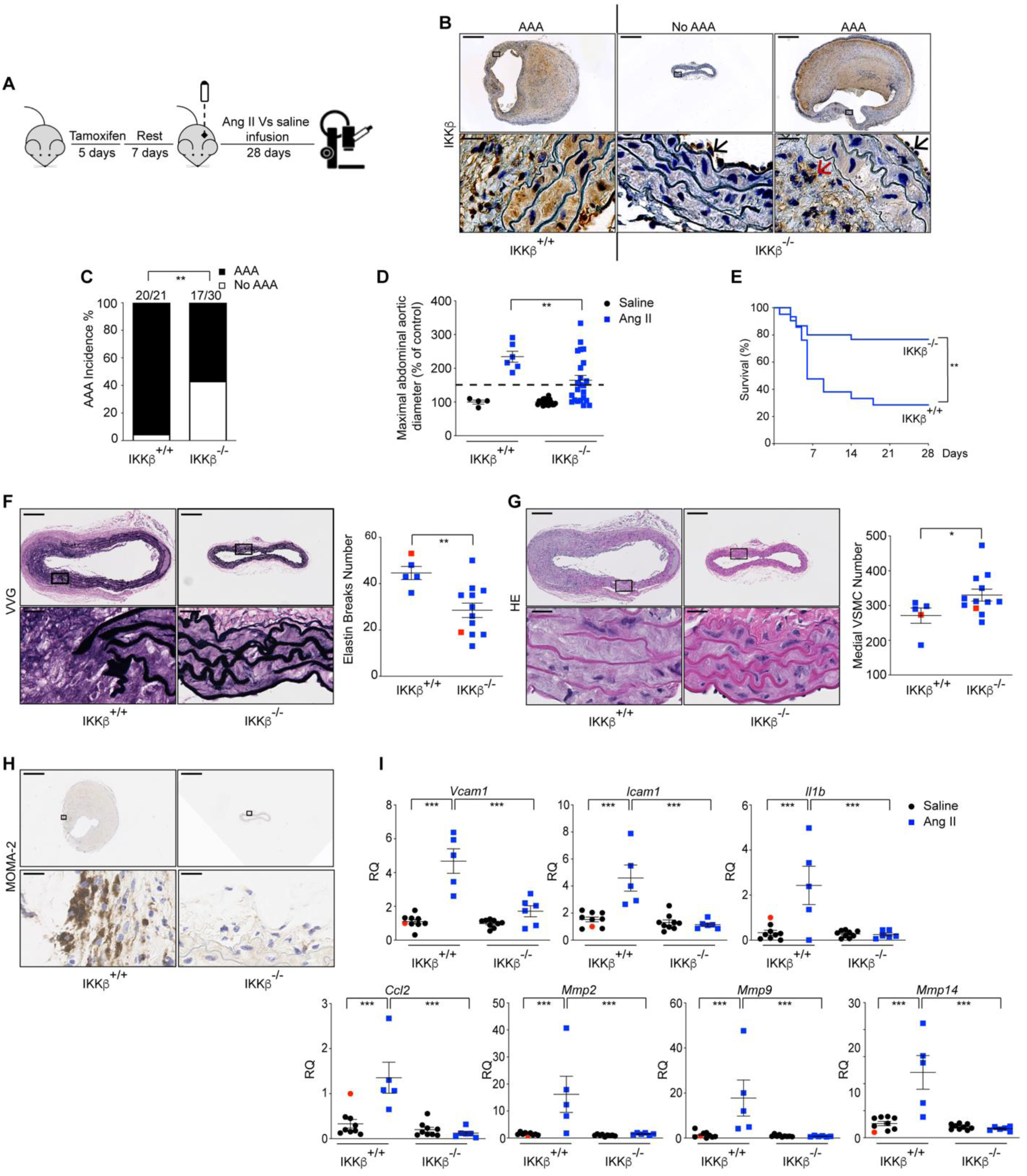
: IKKβ deletion in VSMC prevents AAA development and fatal ruptures. A) Schematic representation of the experimental procedure. Eight weeks-old SMA-CreER^T2^ IKKβ^Flox/Flox^ *ApoE*^-/-^ (IKKβ^-/-^) and their counterpart SMA-CreER^T2^ IKKβ^WT/WT^ *ApoE*^-/-^ (IKKβ^+/+^) male mice were injected with tamoxifen for 5 consecutive days. After 7 days of rest, mice were subjected to saline or Ang II infusion at 1000 ng/kg/min for 28 days. B) Representative histochemistry of conditional deletion of IKKβ in VSMC of SRA sections from IKKβ^+/+^ (*n* = 2) and IKKβ^-/-^ (*n* = 4) mice infused with Ang II. Black and red arrows showing intimal and adventitial staining, respectively, to demonstrate specific IKKβ knockout in the media. Scale bars: 600 μm; Magnification scale bars: 37.5 μm. C) AAA incidence in mice from both groups infused with Ang II at protocol endpoint. ** P < 0.01, Fisher’s exact test. D) Maximal abdominal aortic diameter in mice from both groups infused with saline or Ang II at protocol endpoint. Data were quantified and represented as mean ± SEM. * P < 0.05, 2-way ANOVA with Tukey’s multiple-comparison test. E) Survival curve of IKKβ^+/+^ (*n* = 21) and IKKβ^-/-^ (*n* = 30) mice infused with Ang II for 28 days. *** P < 0.001, Log-rank (Mantel Cox) test. F) Representative Verhoeff–Van Gieson (VVG) stains of SRA sections from IKKβ^+/+^ (*n* = 5) and IKKβ^-/-^ (*n* = 12) mice infused with Ang II and elastin breaks number. Scale bars: 150 μm; Magnification scale bars: 37.5 μm. Red squares indicate the representative stain. Data were quantified and represented as mean ± SEM. ** P < 0.01, 2-tailed Student’s *t* test. G) Representative haematoxylin and eosin (HE) stains of SRA sections from IKKβ^+/+^ (*n* = 5) and IKKβ^-/-^ (*n* = 12) mice infused with Ang II and medial VSMC number. Scale bars: 150 μm; Magnification scale bars: 37.5 μm. Red squares indicate the representative stain. Data were quantified and represented as mean ± SEM. * P < 0.05, 2-tailed Student’s *t* test. H) Representative MOMA-2 histochemistry of SRA sections from IKKβ^+/+^ (*n* = 5) and IKKβ^-/-^ (*n* = 12) mice infused with Ang II. Scale bars: 600 μm; Magnification scale bars: 37.5 μm. I) qPCR analysis of various mRNA expression in TA/SRA lysates from IKKβ^+/+^ (*n* = 9) and IKKβ^-/-^ (*n* = 9) mice infused with saline or IKKβ^+/+^ (*n* = 5) and IKKβ^-/-^ (*n* = 6) mice infused with Ang II at protocol endpoint. Red circles represent the reference. Data were quantified and represented as mean ± SEM. ** P < 0.01, *** P < 0.001, **** P < 0.0001, 2-way ANOVA with Tukey’s multiple-comparison test. RQ: Relative quantification

Although the absence of IKKβ in the vascular wall diminished the development of AAA and fatal ruptures over 28 days of Ang II infusion, up to 57% of the animals developed complex AAA lesions (**Figure 2C** and **Suppl. Figure 1G**). Thus, we next tested if the use of a lower dose of Ang II in our mouse model would show the absolute requirement of IKKβ expressed in VSMC for the development of AAA and its complications. It is noteworthy to mention that animals lacking the expression of IKKβ in the vascular wall and infused with 500 ng/min/kg of Ang II for 28 days did not show any sign of pseudoaneurysm compared to the control group where 40% (6/15) of the IKKβ^+/+^ mice developed AAA associated with 20% of Type I lesions (dilated lumen in the supra-renal region of the aorta with no thrombus), 13% of Type IV lesions (a form in which there are multiple aneurysms containing thrombus, some overlapping, in the SRA area of the aorta) and 7% (1/15) of mortality due to fatal abdominal rupture (**Suppl. Figure 1G and Suppl. Figure 2A-C**). Preservation of both the elastin network and the multilayered medial SMC, reduced MMP2 protein in the media of SRA segments, and a significative inhibitory effect over selected inflammatory transcripts in aorta homogenates paralleled the absence of aortopathy (**Suppl. Figure 2D-G)**. The levels of circulating neutrophils/macrophages chemokines known to be present in human AAA and murine models were also significantly reduced in the VSMC-specific *Ikbkb*-knockout mice (**Suppl. Figure 2H)**. Taken together, the results suggest that the inflammatory function of IKKβ within the vascular wall is required for the recruitment of immune cells, production of MMPs by infiltrating leukocytes and resident VSMC, and the breakdown of the elastin network which ultimately leads to vascular wall remodeling events that are associated with the presence of pseudoaneurysms and ruptures in this model ^39,40^.

### IKKβ governs VSMC switching

The failure of lipid-lowering, anti-hypertensive, anti-inflammatory and anti-protease therapies (indomethacin, fenofibrate, IL-1β neutralization, mast cell inhibition, statins, ACE-inhibitors, β-blockers doxycycline) to reduce AAA progression in humans ^31^ suggest the participation of yet undiscovered or underestimated essential elements, such as VSMC phenotypic switching, that likely remains unresponsive to the aforementioned therapies. In experimental models, Ang II triggers VSMC phenotypic switching through, in part, mTORC1-dependent hypertrophic and anti-autophagic effects, cellular responses that were recently documented as processes leading to early aortopathy in *ApoE*-deficient mice ^39,41,42^. As we previously showed a role of IKKβ in mTORC1-dependent hypertrophic phenotypic response of VSMC ^21^, we next thought to verify the expression of synthetic and contractile markers in aortic tissues derived from IKKβ^+/+^ and IKKβ^-/-^ animals exposed to Ang II for different periods of time. Aortas isolated from animals infused with Ang II for 28 days showed an induced expression of VSMC synthetic markers transcripts involved in collagen expression and remodeling (maturation and turnover) which included the collagen type I alpha 2 chain (*Col1a2*) and the collagen type III alpha 1 chain (*Col3a1*), the prolyl-4-hydroxylase 2 (*P4ha2*) and 3 (*P4ha3*), the lysyl oxidase *(Lox*), the lox like 1 (*Loxl1*), 2 (*Loxl2*) and 4 (*Loxl4*), the MMPs inhibitor TIMP metallopeptidase inhibitor 1 (*Timp1*) as well as the fibrotic/protective marker transforming growth factor beta 1 (*Tgfb1*). The ablation of IKKβ in the vascular wall totally blunted their expression (**Figure 3A**). Remarkably, the expression of the key initiating transcription factor governing this phenotypic switching, *Klf4*, was significantly reduced in IKKβ^-/-^ littermates exposed to Ang II (**Figure 3B**). Using SRA regions isolated from animals exposed for 24 hours to Ang II, we further observed that the preservation of a contractile phenotype is a very early process in our model lacking IKKβ where the relative expression of classical contractile markers myosine heavy chain 11 (*Myh11*) and transgelin (*Tagln*) and the protective autophagic marker autophagy related 5 (*Atg5*) ^42^ were significantly increased after 24 hours of Ang II infusion in a situation where the expression of the inflammatory marker *Vcam1* is not yet modulated (**Figure 3C**). This early preservation of the contractile phenotype is further substantiated by a significantly reduced expression of *Klf4* in IKKβ^-/-^ infused animals littermates exposed to Ang II for 48 hours (**Figure 3D**).

**Figure 3.**
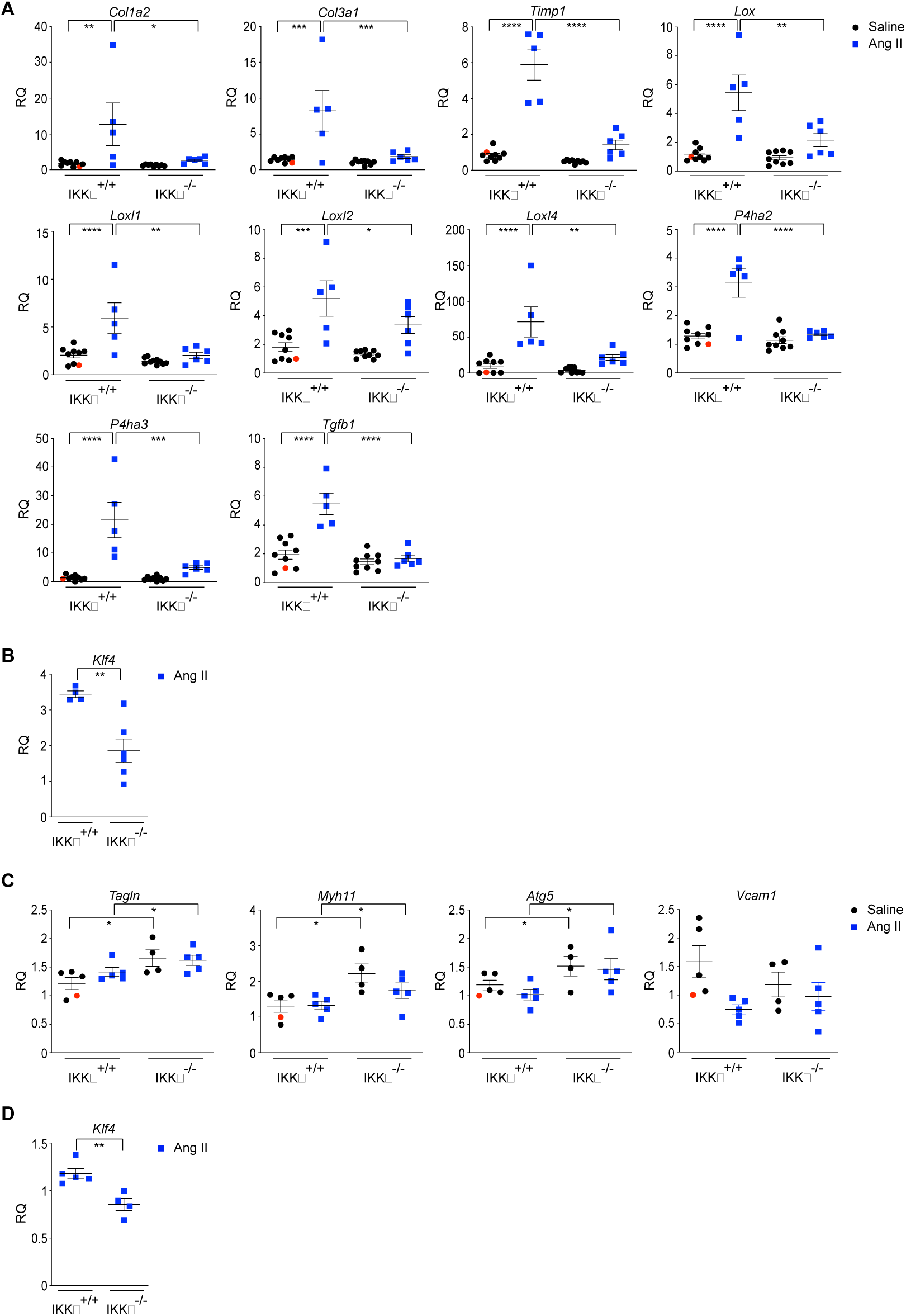
: IKKβ is required for VSMC phenotypic switching. A) qPCR analysis of various mRNA expression in TA/SRA lysates from IKKβ^+/+^ (*n* = 9) and IKKβ^-/-^ (*n* = 9) mice infused with saline or IKKβ^+/+^ (*n* = 5) and IKKβ^-/-^ (*n* = 6) mice infused with Ang II 1000 ng/kg/min for 28 days. Red circles represent the reference. Data were quantified and represented as mean ± SEM. ** P < 0.01, *** P < 0.001, **** P < 0.0001, 2-way ANOVA with Tukey’s multiple-comparison test. RQ : Relative quantification B) qPCR analysis of *Klf4* expression in TA/SRA lysates from IKKβ^+/+^ (*n* = 4) and IKKβ^-/-^ (*n* = 6) mice infused with Ang II 1000 ng/kg/min for 28 days. Data were quantified and represented as mean ± SEM. ** P < 0.01, 2-tailed Student’s *t* test. RQ : Relative quantification C) qPCR analysis of various mRNA expression in SRA lysates from IKKβ^+/+^ (*n* = 5) and IKKβ^-/-^ (*n* = 4) mice infused with saline at day 0 or IKKβ ^+/+^ (*n* = 5) and IKKβ ^-/-^ (*n* = 5) mice infused with Ang II 1000 ng/kg/min for 1 day. Red circles represent the reference. Data were quantified and represented as mean ± SEM. * P < 0.05, 2-way ANOVA with Tukey’s multiple-comparison test. RQ : Relative quantification D) qPCR analysis of *Klf4* expression in SRA lysates from IKKβ^+/+^ (*n* = 5) and IKKβ^-/-^ (*n* = 5) mice infused with Ang II 1000 ng/kg/min for 2 days. Data were quantified and represented as mean ± SEM. ** P < 0.01, 2-tailed Student’s *t* test. RQ : Relative quantification

### IKKβ acts as a VSMC fate-shaping regulator and controls the development of early aortic pseudoaneurysms

VSMC dedifferentiate and initiate the expression of macrophage markers upon cholesterol exposure ^43^. Given the importance of the early sterol biogenesis signature (**Figure 1E and Suppl. Figure S1A**) and the roles of mTORC1 and KLF4 in the rapid polarization of VSMC towards phagocytic-like cells in aortic diseases including dissections and ruptures ^42,44–46^, we next addressed the contribution of IKKβ in the development of early aortic lesions and the presence of degradative SMC phenotype. As shown in **Figure 4A**, from day 2 to day 8 of Ang II infusion, no significant aortic enlargements were noted. However, the absence of IKKβ in the vascular wall led to a reduction in the occurrences of pseudoaneurysms and ruptures (**Table I**), effects that correlated with a diminution in the SMC expressing the macrophage marker galectin 3 (Gal3) (**Figure 4B**). We thus hypothesized that IKKβ was responsible for the early anabolic transcriptomic signature observed in wild-type mice in response to Ang II (**Figure 1E**). To decipher the role of IKKβ in this response, we first performed GSEA analyses in IKKβ^+/+^ and IKKβ^-/-^ mice and generated networks of enriched gene sets. Strikingly, these analyses show that anabolic processes linked to ribosome biogenesis, mRNA translation and mitochondrial respiration are not induced in IKKβ^-/-^ mice in response to Ang II (**Figure 4C**). However, knock-out animals still respond to Ang II, as shown by volcano plots and heatmap representation of DET (**Suppl. Figure 3A-B**), and the induction of genes linked to lipid catabolism (**Figure 4C**). This suggests that IKKβ has a profound implication in vascular cells and determines the metabolic fate in response to Ang II. Further analyses of the most induced transcripts confirm that the individual transcripts associated with oxidative phosphorylation (OXPHOS), ribosome biogenesis, sterol biosynthesis, UPR and fibrosis are not induced or less induced in IKKβ^-/-^ mice in response to early Ang II signaling, and that the lipid catabolism signature is specific to IKKβ^-/-^ mice (**Figure 4D**). Importantly, the anabolic signature of IKKβ^+/+^ mice is strongly associated with mTORC1, a main regulator of these cellular processes ^10^.

**Table 1.** AAA classification in both groups infused with saline or Ang II 1000 ng/kg/min at different endpoint (n = 6)

| Groups | Type | Day 0 | Day 2 | Day 3 | Day 4 | Day 8 |
| --- | --- | --- | --- | --- | --- | --- |
| IKKβ<sup>+/+</sup> | 0 | 100% | 50% | 33% | 0% | 17% |
| IKKβ<sup>-/-</sup> | | 100% | 100% | 83% | 83% | 50% |
| IKKβ<sup>+/+</sup> | I | 0% | 0% | 0% | 0% | 0% |
| IKKβ<sup>-/-</sup> | | 0% | 0% | 0% | 0% | 0% |
| IKKβ<sup>+/+</sup> | II | 0% | 17% | 33% | 50% | 50% |
| IKKβ<sup>-/-</sup> | | 0% | 0% | 17% | 17% | 33% |
| IKKβ<sup>+/+</sup> | III | 0% | 0% | 0% | 0% | 0% |
| IKKβ<sup>-/-</sup> | | 0% | 0% | 0% | 0% | 0% |
| IKKβ<sup>+/+</sup> | IV | 0% | 33% | 0% | 33% | 17% |
| IKKβ<sup>-/-</sup> | | 0% | 0% | 0% | 0% | 0% |
| IKKβ<sup>+/+</sup> | Rupture | 0% | 0% | 33% | 17% | 17% |
| IKKβ<sup>-/-</sup> | | 0% | 0% | 0% | 0% | 17% |

**Figure 4.**
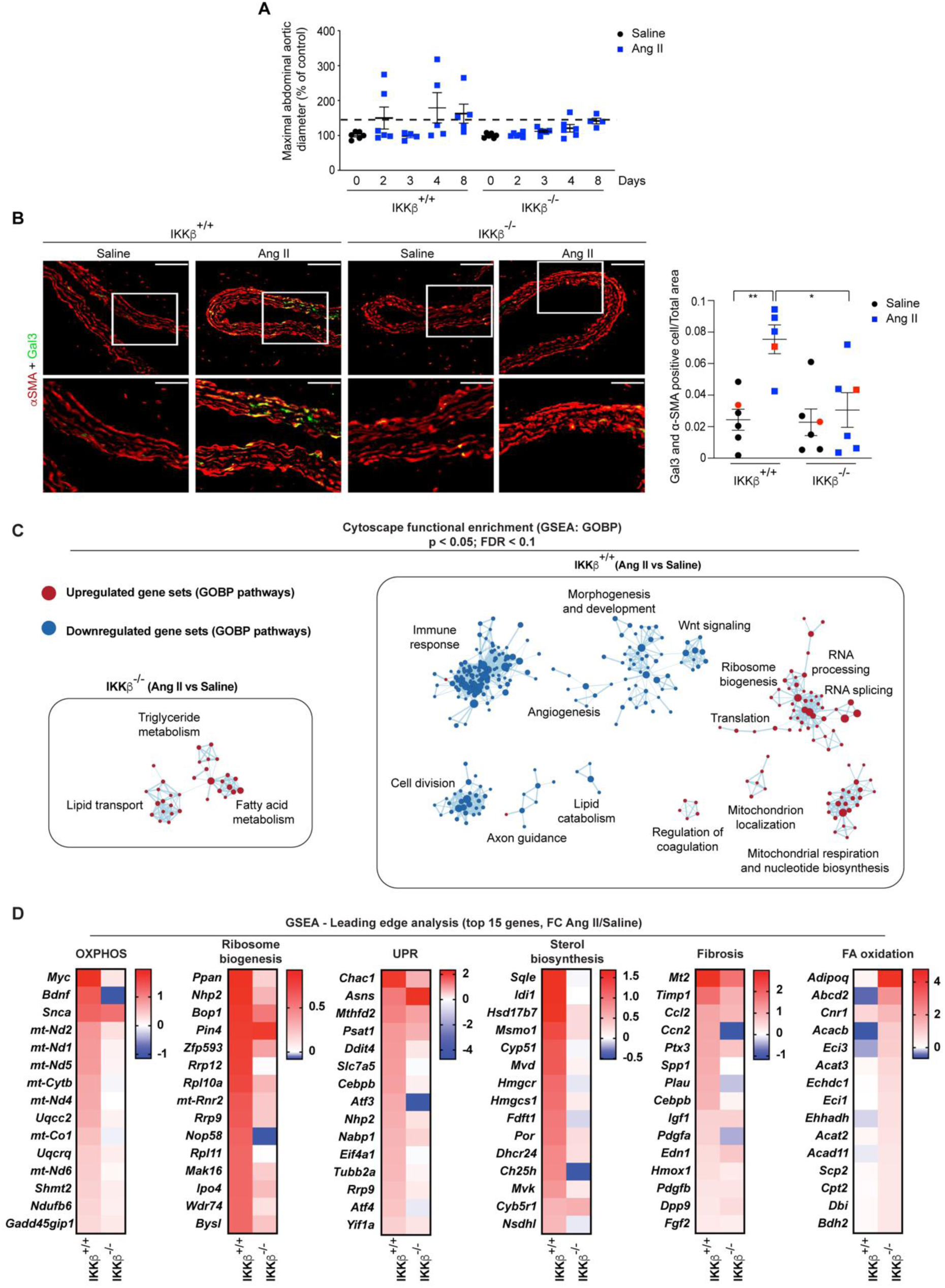
: IKKβ drives VSMC toward a degradative phenotype. A) Maximal abdominal aortic diameter in mice from both groups infused with saline or Ang II 1000 ng/kg/min at different endpoint. Data were quantified and represented as mean ± SEM. B) Representative α-SMA and Gal3 immunofluorescent costaining of SRA sections from IKKβ^+/+^ (*n* = 6) and IKKβ^-/-^ (*n* = 6) mice infused with saline at day 0 or IKKβ^+/+^ (*n* = 5) and IKKβ^-/-^ (*n* = 6) mice infused with Ang II 1000 ng/kg/min for 4 days and quantification. Scale bars: 40 μm; Magnification scale bars: 20 μm. Red squares indicate the representative stain. Data were quantified and represented as mean ± SEM. * P < 0.05, ** P < 0.01, 2-way ANOVA with Tukey’s multiple-comparison test. C) Cytoscape network representation of enriched gene sets in IKKβ^+/+^ and IKKβ^-/-^ mice in response to early Ang II signaling. Nodes represent gene sets, node size represents the number of genes within gene sets, and edges indicate the similarity between gene sets. Red indicates upregulated gene sets, and blue indicates downregulated gene sets. D) Heatmap representations of transcript fold change (Log2) in response to Ang II treatment (versus saline) for selected GSEA pathways. Transcripts are ranked according to fold change from highest (red) to lowest (blue).

To further substantiate the observed fate-shaping properties of IKKβ, we next address the question in cultured VSMC. In serum-starved VSMC, we recapitulate the inhibitory effect of the selective IKKβ inhibitor MLN120b on the phosphorylation of the mTORC1 substrate ribosomal protein S6 kinase (p70S6K) in Ang II-treated cells ^21^ (**Figure 5A**). However, these short kinetics cannot recapitulate and capture the transcriptional-dependent fate-shaping processes occurring in these cells. As cultured VSMC rapidly adopt a synthetic phenotype once adhered to plastic dishes and exposed to serum, we thus exposed them to MLN120b for 6 days and looked at the steady state expression levels of KLF4 and β-catenin, a recently described transcriptional coactivator involved in the polarization of VSMC toward a degradative “macrophage” phenotype ^45^. The use of MLN120b reduced the expression of both phenotypic markers in a concentration-dependant manner (**Figure 5B)**. Furthermore, inhibiting IKKβ also negatively impacted the expression of cellular communication network factor 2 (CCN2; also known as CTGF) *in vitro* and *in vivo* (**Figure 5C-D**), a matricellular protein linked to various fibrotic processes and recently proposed to decrease VSMC adhesion to structural ECM, promoting aortic dissections ^39^. Altogether our results identify IKKβ as a novel VSMC fate-shaping regulator controlling early aortic remodeling events leading to pseudoaneurysms and ruptures.

**Figure 5.**
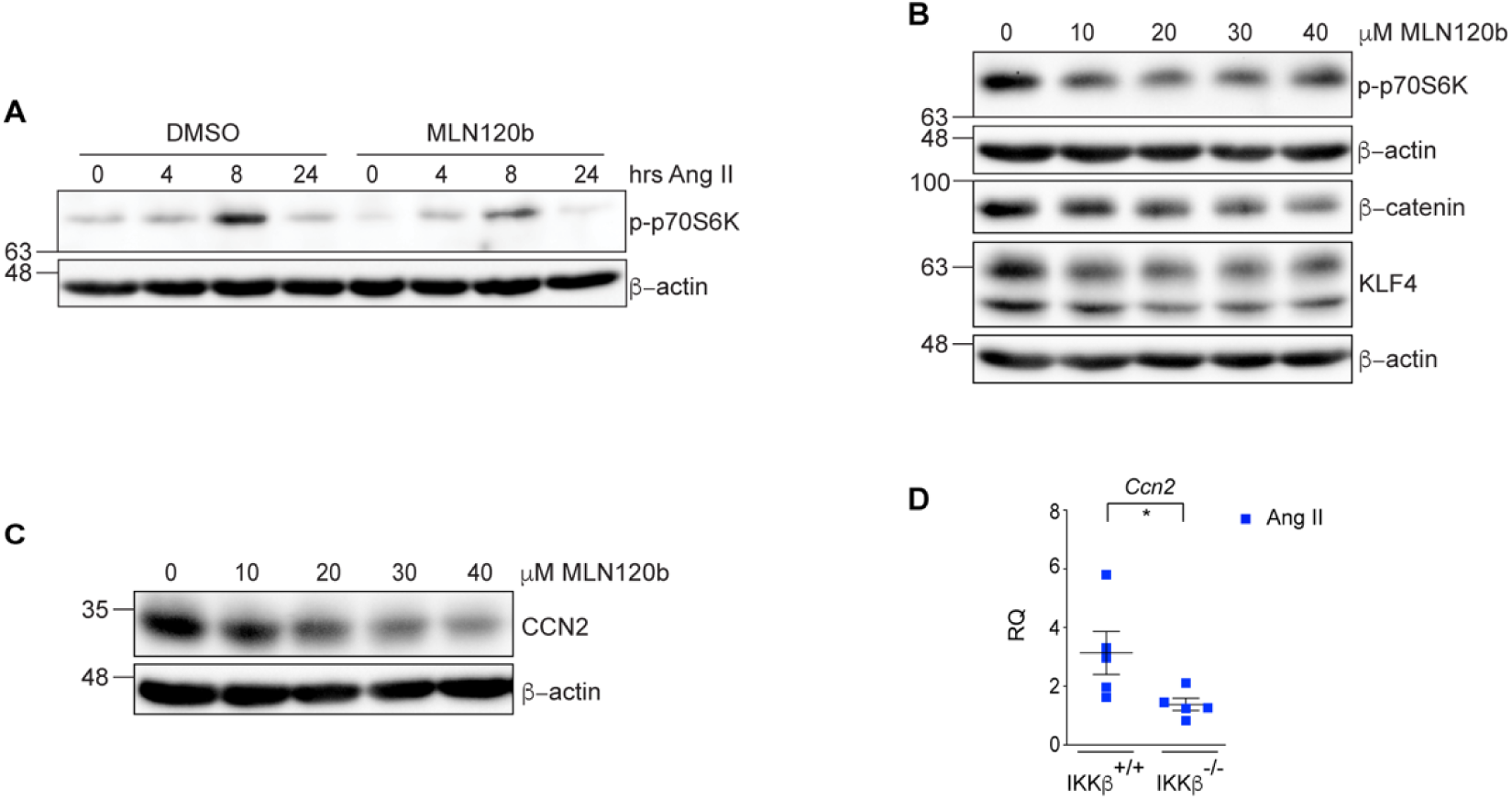
: Fate-shaping properties of IKKβ in cultured VSMC. A) Quiescent VSMC were pre-treated with DMSO or MLN120b (10 µM) for 30 min before Ang II (100 nM) addition for the indicated times. Cell extracts were subject to immunoblotting with the indicated antibodies. Data are representative of 3 independent experiments. B-C) VSMC were treated with either DMSO or the indicated concentrations of MLN120b for 6 days. Cell extracts were subject to immunoblotting with the indicated antibodies. Data are representative of minimally 2 independent experiments. D) qPCR analysis of *Ccn2* expression in SRA lysates from IKKβ^+/+^ (*n* = 5) and IKKβ^-/-^ (*n* = 5) mice infused with Ang II 1000 ng/kg/min for 1 day. Data were quantified and represented as mean ± SEM. * P < 0.05, 2-tailed Student’s *t* test. RQ : Relative quantification

## DISCUSSION

In the present study, we first demonstrate that transcriptomic changes associated with long-term detrimental outcomes of AAA are established as early as 24 hours following Ang II infusion. Moreover, we show that the earliest events underlying the pathophysiology of AAA involve activating anabolic pathways rather than inflammatory ones. Furthermore, in addition to its direct activation of the mTORC1 signaling cascade^21^ and modulating the early anabolic events, IKKβ has a crucial role in the phenotypic switching of VSMC towards a degradative phenotype. Collectively, our study provides a novel regulatory signaling IKKβ-mTORC1-axis in the development and progression of AAA and that the control of the anabolic response through IKKβ deserves considerations in the design of future studies aiming at the discovery and characterization of potential therapeutic targets for the treatment of early AAA lesions. From a clinical perspective, it has been suggested that there may be an association between anabolic events and aortic diseases. Notably, anabolic reactions have been observed to elevate with advancing age ^10^, which represents the predominant clinical characteristic of AAA. Furthermore, recent case reports have established a potential correlation between the usage of anabolic steroids and the occurrence of thoracoabdominal aneurysms and aortic dissection ^47,48^.

For more than two decades, studies used Ang II infusion in hyperlipidemic mice models to gain mechanistic insight into human AAA ^34^. In this context, Ang II is considered the primary inflammatory trigger leading to a complex response where cytokines and MMPs produced by resident fibroblasts, VSMC, endothelial cells, and innate/adaptative immune cells lead to the deleterious remodeling of the vascular wall^49^. In addition to this inflammatory component, accumulating evidence indicates that VSMC within remodeled vascular regions exhibit notable flexibility and can go under various phenotypes with potential clinical implications, which can be beneficial or detrimental^50^. Understanding the distinct stages and mechanisms involved in VSMC activation, such as transitioning from quiescence to invasion of remodeled vascular areas and fate diversification, is crucial for devising therapeutic interventions aimed at mitigating pathological VSMC behaviors while preserving or enhancing their stabilizing functions. In this study, using a highly characterized Ang II-infused *ApoE*-deficient mice model, we discovered an early molecular anabolic signature associated with activating the mTORC1 pathway (**Figure 1**). Dysregulation in mTORC1 activation was recently shown to promote aortopathy by inducing a phenotypic switching of VSMC to a degradative phenotype thus inducing proteolysis, extracellular matrix degeneration, and inflammation in the aortic wall ^45^. While blocking mTOR signaling has shown to be a promising therapeutic target to limit both thoracic aortic aneurysm and AAA development ^51–53^, a recent study indicated that targeting mTORC1 by using rapamycin promotes aortic dissection ^39^. Interestingly, this was associated with the continuous and mTORC1-independent induction of CCN2 by Ang II, decreasing VSMC adhesion to structural ECM and allowing extensive dissection ^39^. In the current study, we show that targeting upstream of mTORC1 (i.e. conditional deletion of IKKβ in VSMC) not only reduces the aortic dilatation but also dramatically decreases the aortic dissection and fatal ruptures. Based on our detailed analysis of the SRA segments in the early (one to four days) or late (28 days) infusion protocols, we propose that IKKβ plays pivotal roles in the initiation and disease progression through at least three disctict mechanisms: 1) through its ability in controlling mTORC1 in VSMC exposed to Ang II, IKKβ rapidly engages anabolic events leading to A) the generation of magrophage-like VSMC responsible for early vascular wall remodeling events as well as B) a rapid metabolomic reprogramming toward OXPHOS; 2) combine with increased ribosome synthesis and the UPR, the above events create the proper conditions favoring ROS production ^54^ and exacerbated inflammatory response in the vascular wall, leading to 3) leucocytes infiltration and sustained NF-κB-dependant induction of *Ccn2* ^55^ culminating to aortic dissections and fatal ruptures. Rapid monocytes recruitment ^32^ and/or the presence of sparse tissue-resident macrophages could play a role in the rapid vascular wall remodeling. However, the early occurrence of the mTORC1 signature (24 hours) along with the VSMC switching towards degradative phenotype suggests a direct role of IKKβ in these effector cells. Importantly, IKKβ might also have other roles in VSMC and macrophages as recently demonstrated ^8,56^.

Through the pleiade of substrates including SREBP, eIF4E-binding protein 1/2 (4E-BP1/2), ribosomal protein S6 kinases 1/2 (p70S6K1/2), the Ser/Thr kinase ULK1, the transcriptional co-activator PGC-1α and the transcription factor YY-1, the effector anabolic kinase complex mTORC1 controls the genesis of several essential macromolecules (lipids, proteins, ribosomal RNAs) and directly influence cellular energy metabolism by either inhibiting autophagy and/or increasing mitochondria functions/biogenesis^10^. Although essential for cell growth, mTORC1 activation is also linked to aging (decrease longevity) ^10^ and fibrosis ^38^, biological processes that play major roles in developing AAA. We propose here that the dramatic reduction of aortic dissections and fatal ruptures observed in the IKKβ^-/-^ animals are explained by the fact that IKKβ acts has a molecular hub in the vascular wall, allowing vasoactive molecules such as Ang II to control both the classical inflammatory effectors such as NF-kB and mTORC1. In this context, it would be interesting to verify if the failure of the vasoactive agent norepinephrine to promote AAA is related to its incapacity of stimulating the IKKβ-mTORC1-dependent anabolic/inflammatory responses ^57^.

In conclusion, our study identified the previously unrecognized role of IKKβ in AAA development and progression and a novel therapeutic target. However, it is important to emphasize that prolonged systemic IKKβ inhibition has context-dependent effects on the immune system and inflammatory processes ^58^. Therefore, inhibiting IKKβ locally or systemically in humans in the context of AAA requires further investigations.

## ACKNOWLEDGMENTS

We thank Dr. Hélène Girouard (Université de Montréal) for sharing her expertise and equipments for blood pressure monitoring. We are also grateful to Etienne Durette and Dr. Wendy van Zuijlen for data collection and the establishment of the experimental animal models.

## FUNDING

This work was supported by research grants from the Canadian Institutes of Health Research (CIHR; MOP-123482) and the Heart and Stroke Foundation of Canada (HSFC; G-16-00014208) to M.J.S.

## DISCLOSURE

None

## SUPPLEMENTAL MATERIAL

Expanded Methods

Tables S1 and S2

Figure S1-S3

REF #59 to #73

## NON-STANDARD ABBREVIATIONS AND ACRONYMS

AAA: abdominal aortic aneurysm
ECM: extracellular matrix
VSMC: vascular smooth muscle cells
IKKβ: inhibitor of nuclear factor kappa B kinase subunit beta
NF-kB: nuclear factor kappa B
Ang II: angiotensin II
ApoE: apolipoprotein E
SRA: suprarenal
RNA-seq: RNA sequencing
DET: differentially expressed transcripts
GSEA: Gene Set Enrichment Analyses
mTORC1: mammalian target of rapamycin complex 1
UPR: unfolded protein response
TFBS: transcription factor binding sites
KLF: Krüppel-like factors
MMP: matrix metallopeptidase
CCN2: cellular communication network factor 2

## Notes

### Competing Interest Statement

The authors have declared no competing interest.

